# New efficient intercellular spread mode of respiratory syncytial virus contributes to neutralization escape and persistence

**DOI:** 10.1101/2022.02.01.478688

**Authors:** Wei Zhang, Xue Lin, Lu-Jing Zhang, Li Chen, Yong-Peng Sun, Jun-Yu Si, Min Zhao, Guang-Hua Wu, Lu-Ting Zhan, Ying-Bin Wang, Ning-Shao Xia, Zi-Zheng Zheng

## Abstract

There is no licensed vaccine or therapeutic antibody for respiratory syncytial virus (RSV). The induction of high-titer, potent neutralizing antibodies cannot completely inhibit breakthrough infection and enhanced respiratory disease (ERD), encouraging a focus on the relationship between virus intercellular spread and neutralizing antibodies. By blocking the known intercellular spread modes and with the aid of ultrahigh-resolution imaging, we observed a new efficient mode of intercellular spread in which RSV-infected cells directly transfer viral materials (including viral replication factories) to neighboring cells through protruding open-ended microfilament-rich intercellular nanotubes. The new mode is virion-independent and antibody-insensitive, beginning as early as 3 h post infection. Furthermore, replication-defective viral genomes (DVGs) might also utilize the new mode, facilitating the establishment of latent viral infections. Therefore, our data provide a new perspective on RSV cell-to-cell spread and might help to explain the immune escape and latent persistence of paramyxoviruses.

## Introduction

Human respiratory syncytial virus (RSV) is the leading cause of hospitalization in children with acute lower respiratory tract infection (ALRI) and the most important viral cause of ALRI mortality in childhood (Nair et al., 2010). The RSV F and G proteins are the two main neutralizing antigens and play key roles in virus entry and attachment, respectively (McLellan et al., 2013c). RSV infects human respiratory tract epithelial cells (Hall, 2001). In vitro, cells infected with RSV form syncytia and filamentous surface projections presenting F and G glycoproteins (Collins et al., 2013a). Similar to those of other paramyxoviruses, the RSV replication process is characterized by the formation of inclusion bodies (IBs) (Norrby et al., 1970), which consist of the nucleoprotein (N), the phosphoprotein (P), the large polymerase subunit (L), and the transcription processivity factor (M2-1), as well as viral genomic RNA and messenger RNA (mRNA) (Rincheval et al., 2017). IBs are believed to regulate MDA5-related innate immune responses (Lifland et al., 2012) and are viral factories where viral RNA synthesis occurs (Rincheval et al., 2017, Santangelo et al., 2006).

To date, there is no licensed vaccine for RSV, and there is only one prophylactic antibody, palivizumab (1998), although the virus was isolated nearly 70 years ago. The development of an RSV vaccine is considered one of the biggest challenges in vaccinology given the risk of vaccine-induced enhanced respiratory disease (ERD). The discovery of the pre-F conformation has provided opportunities and insights for the development of RSV vaccines (McLellan et al., 2013b, McLellan et al., 2013a). However, at suboptimal doses, the recombinant pre-F-based subunit vaccine DS-CAV1 still has a high risk of inducing ERD despite stimulating robust levels of neutralizing antibodies and a strong Th1-biased response (Schneider-Ohrum et al., 2017). Additionally, and interestingly, there is a hook effect regarding the severity of inflammation as the antigen dose decreases and the level of viral replication in the lungs increases. It seems that once antigen-induced immunity fails to prevent the virus from causing a breakthrough infection, viral replication and pathological processes in the lungs run out of control despite the existence of immune memory. The mechanism of this process remains to be elucidated. Therefore, to understand how RSV breakthrough infection occurs in the presence of high titers of neutralizing antibodies, we focused on the cell-to-cell transmission of RSV.

For a long time, our understanding of RSV cell-to-cell transmission has been limited to cell-free virion dissemination and the formation of syncytia with low efficiency. In monolayer cell culture systems, mature progeny virions germinate from the apical surfaces of polarized cells (Santangelo and Bao, 2007), but the majority of them cannot be released from the host cells (Collins et al., 2013a, Huong et al., 2016). In addition, cell-free virions are unstable, fragile and prone to loss of infectivity (Collins et al., 2013a, Liljeroos et al., 2013). Masfique and his colleagues (Mehedi et al., 2016) described another interesting intercellular delivery mode in which RSV-infected cells utilize filopodia to deliver nonshedding virions to neighboring cells. It is worth noting that, similar to the previously reported mode for HIV-1 (Schiffner et al., 2013), this mode is also essentially virion-dependent. Both virion-dependent modes of transmission face some obstacles before they can successfully induce infection.

In this study, we discovered a new mode of cell-to-cell transmission through a series of rigorous experiments based on optical imaging. In this mode, the infected cells became more active in connecting with neighboring cells and delivered viral replication factories (IBs) to target cells via open-ended intercellular nanotubes. This new mode of transmission successfully induced efficient replication of the virus in the target cells at the early stage of infection before virus particle assembly and even transferred the replication-defective viral genomes (DVGs) to adjacent cells, thus promoting the latent persistence of RSV. The results suggest that RSV uses a new mechanism of cell-to-cell transmission, and this work might provide a reference for other viral studies.

## Results

### Neutralizing antibodies inhibited virion-based infection but could not completely block intercellular spread

To understand the possible effects of neutralizing antibodies on cell-to-cell transmission, we tried to simulate different modes. First, for cell-free virion dissemination, HEp-2 cells were infected with RSV-A2-mKate (multiplicity of infection [MOI]=4) and incubated for several hours in the presence of the mAb 5C4 (a pre-F-specific potent neutralizing antibody). Then, a mixture of dissociated mature virions and antibodies was added to the prepared uninfected cells to examine the infectivity of cell-free virions in the presence of antibodies. As shown in Figure 1A-1B, the blockade of cell-free virions with the mAb 5C4 increased with increasing antibody concentration, and the high-titer of antibody was able to completely neutralize cell-free virions generated by 2.5×105 infected cells (20 h post infection [h.p.i.]) in a volume of 2 ml for at least 6 h.

**Figure 1.**
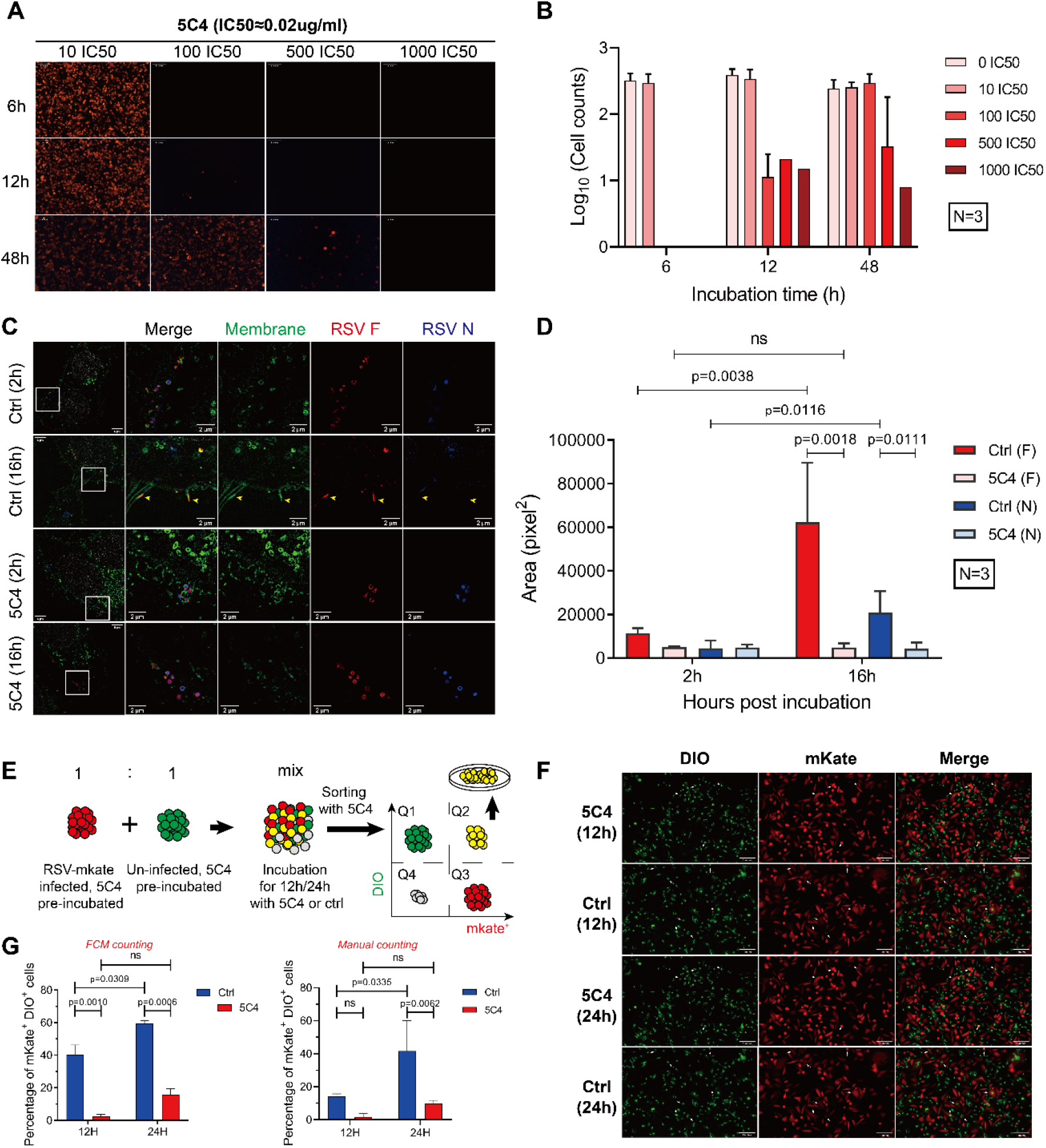
Influence of neutralizing antibodies on intercellular viral spread. (A-B) Inhibitory effects of neutralizing antibodies on cell-free virion infection under appropriate conditions. HEp-2 cells were infected with RSV-A2-mKate and then preincubated with the mAb 5C4 at 20 h.p.i. Then, media containing virions and mAbs were added to uninfected HEp-2 cells to evaluate infectivity. Positive cells were detected by fluorescence microscopy (A) and counted (B). (C-D) Neutralizing antibodies inhibited virion-based cell-to-cell transmission-induced infection. RSV A2-infected HEp-2 cells were prewashed at 20 h.p.i. and then incubated with 100 IC50 of 5C4 or an isotype control. The infected cells were scraped, added to the layer of uninfected cells for a short incubation and finally washed off. Sixteen hours later, the cells were fixed and stained with wheat germ agglutinin (WGA, membrane), motavizumab (RSV F) and 11H8 (RSV N) to examine successful infection. The yellow arrows indicate the budding virions. (E-F) Neutralizing antibodies could not completely inhibit intercellular infection. RSV-A2-mKate-infected cells (donor cells, mKate+DIO-) were preincubated with 100 IC50 of 5C4 and then mixed with uninfected cells (receptor cells, DIO+mKate-) for further incubation in 5C4-containing medium. Finally, the percentage of DIO+mKate+ cells was analyzed via flow cytometry. (E) Sketch of the experimental scheme. (F) Fluorescence microscopic images of mixed cells before flow cytometry; the white arrows indicate DIO+mKate+ cells. (G) Quantitative analysis of the images in (F) and data acquired from flow cytometry. The results are presented as the geometric mean ± SD. The following source data is available for figure 1: **Source data 1.** This spreadsheet contains data used to generate graphs in Figure 1B, 1D, 1G.

Second, we tested whether the filopodia-driven virion intercellular delivery mode, which is virion-dependent, was completely blocked by a proper mAb concentration (as expected). In a simulation model, RSV-A2-infected cells (preincubated with the high-titer mAb 5C4) were scraped with preservation of their filopodia as much as possible and then coincubated with uninfected cells for several hours. The ultrahigh-resolution microscopy data (Figure 1C, 1D) showed that the virions remained in the receptor cell membranes in the presence of antibodies, while the virions successfully caused infection and produced filamentous progeny in the control sample without antibodies. Therefore, the above two virion-based intercellular diffusion modes were completely blocked by high titers of neutralizing antibodies.

Then, we utilized a coculture system to analyze the overall effect of the antibody on RSV intercellular diffusion (Figure 1E-1F). Infected cells (donor, red) were cocultured with uninfected cells (receptor, green) in the presence or absence of the high-titer mAb 5C4. The double-positive cells were then counted. We found that the high-titer mAb 5C4 significantly, but not completely, suppressed intercellular diffusion, implying that there might be an unknown intercellular diffusion mode that is insensitive to high-titer neutralizing antibodies.

### RSV-infected cells developed more open-ended intercellular nanotubes and became more active in contact with neighboring cells

To further analyze the possible spread mode, we observed the characteristics of infected cells in more detail. Viral infection-induced cytoskeletal remodeling has been reported to occur in many viruses (Murti and Goorha, 1983, Bukrinskaya et al., 1998, Igakura et al., 2003, Fantuzzi et al., 2003); this remodeling usually results in changes in cell morphological features and motility. Therefore, we analyze the differences between infected and uninfected HEp-2 cells from several perspectives (Figure 2). At 24 h.p.i., the cells were fixed, permeabilized, and then stained with WGA and 4’,6-diamidino-2-phenylindole (DAPI). Our results showed that infected HEp-2 cells (as well as A549 cells) developed more filopodia-like structures than uninfected cells, consistent with a recent report (Mehedi et al., 2016). However, only open-ended intercellular nanotubes that connected two cells were considered in this study. Aside from the number, the length and width of these intercellular nanotubes were also greater than those of the nanotubes in uninfected cells (Figure 2A, 2B). A wound healing assay was performed to evaluate the influence of RSV infection on cell motility (Figure 2C, 2D). After approximately 12 h of culture, many more cells had moved into the wound area in the infected samples than in the uninfected samples, although the decrease in wound area was slower in the infected samples than in the uninfected samples. This seemingly contradictory result reflects the fact that infected cells move faster than uninfected cells. Furthermore, we tracked the movement of cells; consistently, we found that infected cells moved farther than uninfected cells in the same amount of time (Figure 2—figure supplement 1). The evidence thus far indicated that RSV-infected cells presented more active features, including motility and contact with neighbors, than uninfected cells.

**Figure 2.**
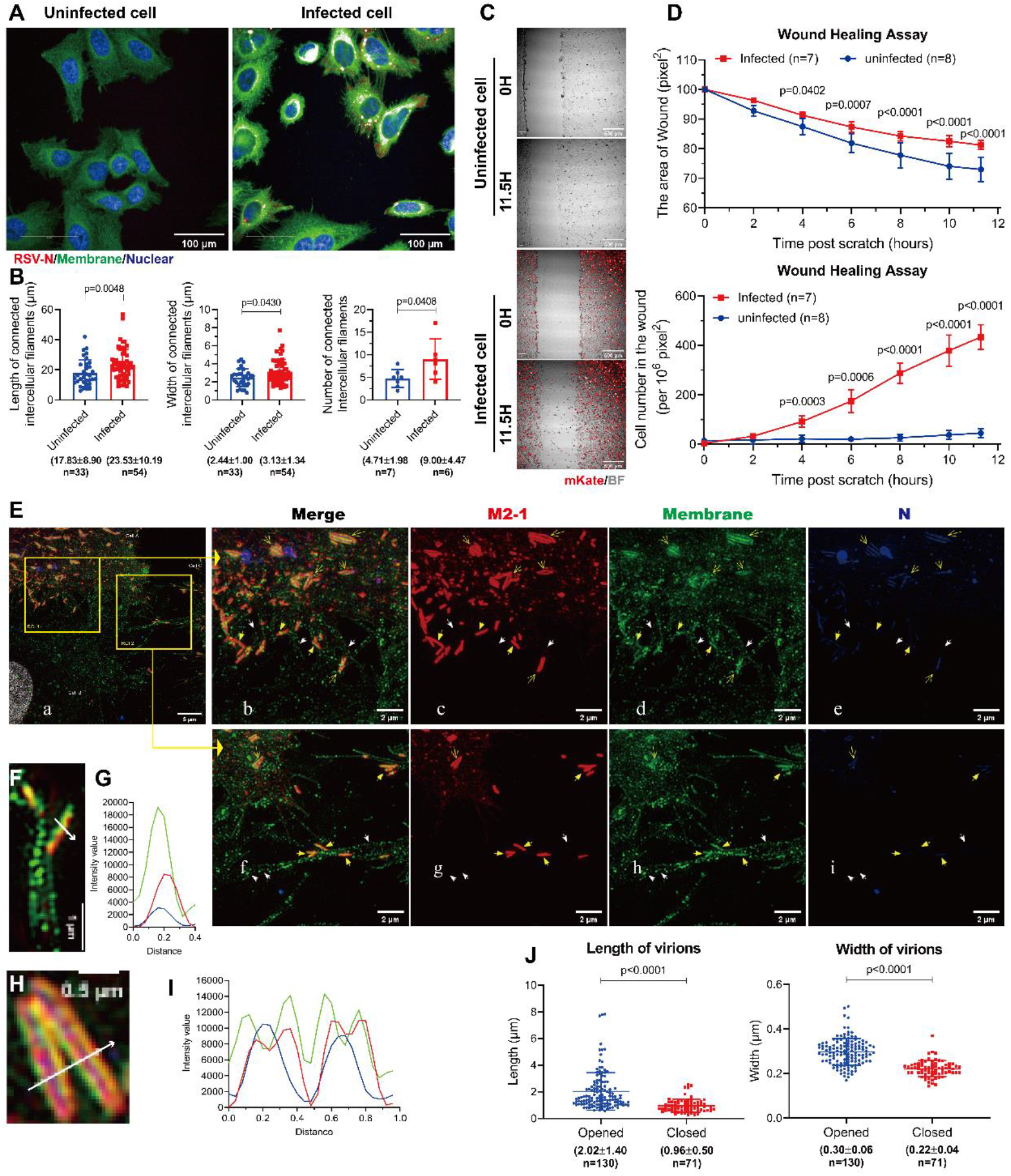
RSV infection increased cell movement and the formation of intercellular nanotubes in which viral proteins were distributed. (A-D) Change in cell morphology induced by RSV infection. (A) Morphological comparison of RSV-infected and uninfected HEp-2 cells. Cells were fixed and stained with DAPI (nuclear), WGA (membrane) and the mAb 11H8 (RSV N protein), and images were acquired with the confocal module of Opera Phenix. (B) Comparison of intercellular connected nanotubes between RSV-infected and uninfected cells. The length, width and number of intercellular nanotubes were examined. (C-D) Wound healing assay. (E-J) Direct migration of RSV viral materials from cell to cell. (E) Observation of intercellular connected nanotubes of RSV-infected cells via ultrahigh-resolution microscopy. RSV-infected cells were fixed and stained with WGA (membrane), the mAb 11H8 (RSV N) and the mAb RSV5H5 (RSV M2-1). Images with ultrahigh resolution were acquired via DeltaVision OMX. The white notched arrows indicate the direct viral material spread in the tunnels of intercellular connected nanotubes (“in the tunnel”). The yellow filled arrows indicate the direct migration of closed virions on the surfaces of intercellular nanotubes (“virion-based cell-to-cell transmission”). The yellow open arrows indicate opened virions. (F-I) Two types of virions and their line profiles. (J) Statistical information of the size parameters of two types of virions that were randomly selected from more than ten images. For the statistical assay, a nonparametric statistical test was performed with GraphPad Prism version 7.00. The results are presented as the arithmetic mean ±SD. The following source data is available for figure 2: **Source data 1.** This spreadsheet contains data used to generate graphs in Figure 2B, 2D, 2G, 2I, 2J. **Figure supplement 1.** RSV infection increased cell movement. **Figure supplement 2.** The morphological features of infected cells under the CLEM. **Figure supplement 3.** The distribution and the abundance of pre-F and post-F on infected cells.

### RSV might utilize a highway to directly transit viral materials (IBs) from cell to cell beginning in the early stage of infection

Ultrahigh-resolution microscopy and correlative light and electron microscopy (CLEM) were utilized to observe the infected cells in more detail (Figure 2E, Figure 2—figure supplement 2). We noted that the viral proteins N and M2-1 were located in open-ended intercellular nanotubes (Figure 2E, S2B, white notched arrows), suggesting the possible transfer of viral materials. In addition, two types of virions were observed on the surfaces of infected cells (Figure 2E, S2A, yellow filled arrows, yellow open arrows); these types differed in the abundance of pre-F (Figure 2—figure supplement 3). Interestingly, on the surfaces of intercellular nanotubes and filopodia, shortened, narrowed and closed virions were distributed, implying that possible viral material transfer might occur in combination with filopodia-driven virion intercellular delivery (Mehedi et al., 2016). The other virions on the basal side of the cell body membrane were lengthened, widened and opened (Figure 2F-2J). These virions might have been more mature.

To further study possible viral material transfer, we needed to separate it from virion-based spread. Fortunately, material transfer, a virion-independent dissemination mode, should theoretically start before virion assembly begins. Hence, we monitored activities in the early period of RSV infection (from 2 h.p.i. to 24 h.p.i.) by staining the N protein to detect IBs and the F protein to detect progeny virion assembly. As expectedly, in the early stage of RSV infection (3 h.p.i.) before virion assembly, viral materials were already detected in the intercellular nanotubes (Figure 3, white notched arrows). Obvious virion assembly on the plasma membrane did not occur until 11 h.p.i. (Figure 3, yellow filled arrows), which was the basis of filopodia-driven virion intercellular delivery. Thus, we determined that the transfer of viral materials through intercellular nanotubes might be the major intercellular spread mechanism in the early stage.

**Figure 3.**
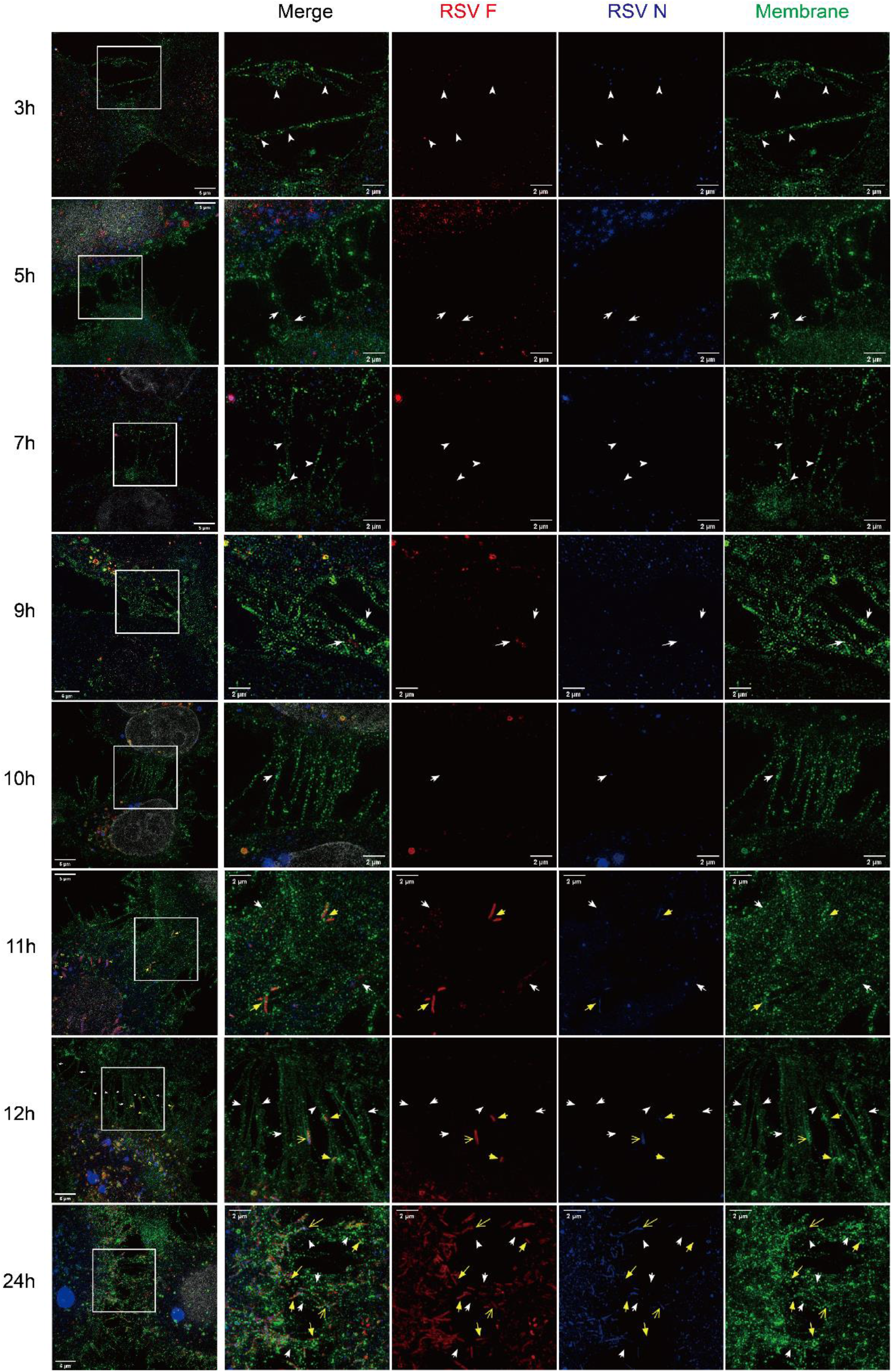
Viral materials were detectable in the tunnels very early before virus budding. HEp-2 cells on slides were infected with RSV-A2 at an MOI of 2, fixed at different timepoints and then stained with WGA (membrane), the mAb 11H8 (RSV N) and motavizumab (RSV F). Images in ultrahigh-resolution mode were acquired via DeltaVision OMX. The white notched arrows indicate viral material spread in the tunnels of intercellular connected nanotubes. The yellow filled arrows indicate the transmission of closed virions on the surfaces of intercellular connected nanotubes. The yellow open arrows indicate opened virions.

To clarify the viral materials in the intercellular nanotubes, we first costained four nucleocapsid/polymerase proteins of RSV (N, P, L and M2-1). Unexpectedly, we found that these four proteins colocalized in the nanotubes (Figure 4A-4C, white notched arrows), which indicated that these proteins might be transported through the tunnels in the form of IBs. We also explored the relative localization of the N protein and RSV RNA in the nanotubes, and the results showed that genomic RNA was always localized with the N protein, while mRNA was probably not (Figure 4D-4E, white notched arrows). The composition of these intercellular nanotubes was determined by costaining the N protein, F-actin and β-tubulin as well as the membrane. Polymerized F-actin (microfilaments) was considered the primary skeleton of these nanotubes; however, β-tubulin was only sporadically present in the tunnels (Figure 4F). Therefore, the intercellular nanotubes referred to above were tunneling nanotube (TNT)-like. In addition, the TNT-like nanotube membranes had a strong WGA signal (Figure 4G-4H); this was a feature of RSV-infected cell membranes due to their rich content of viral transmembrane glycoproteins, suggesting that the source of the nanotube membranes might be infected cells.

**Figure 4.**
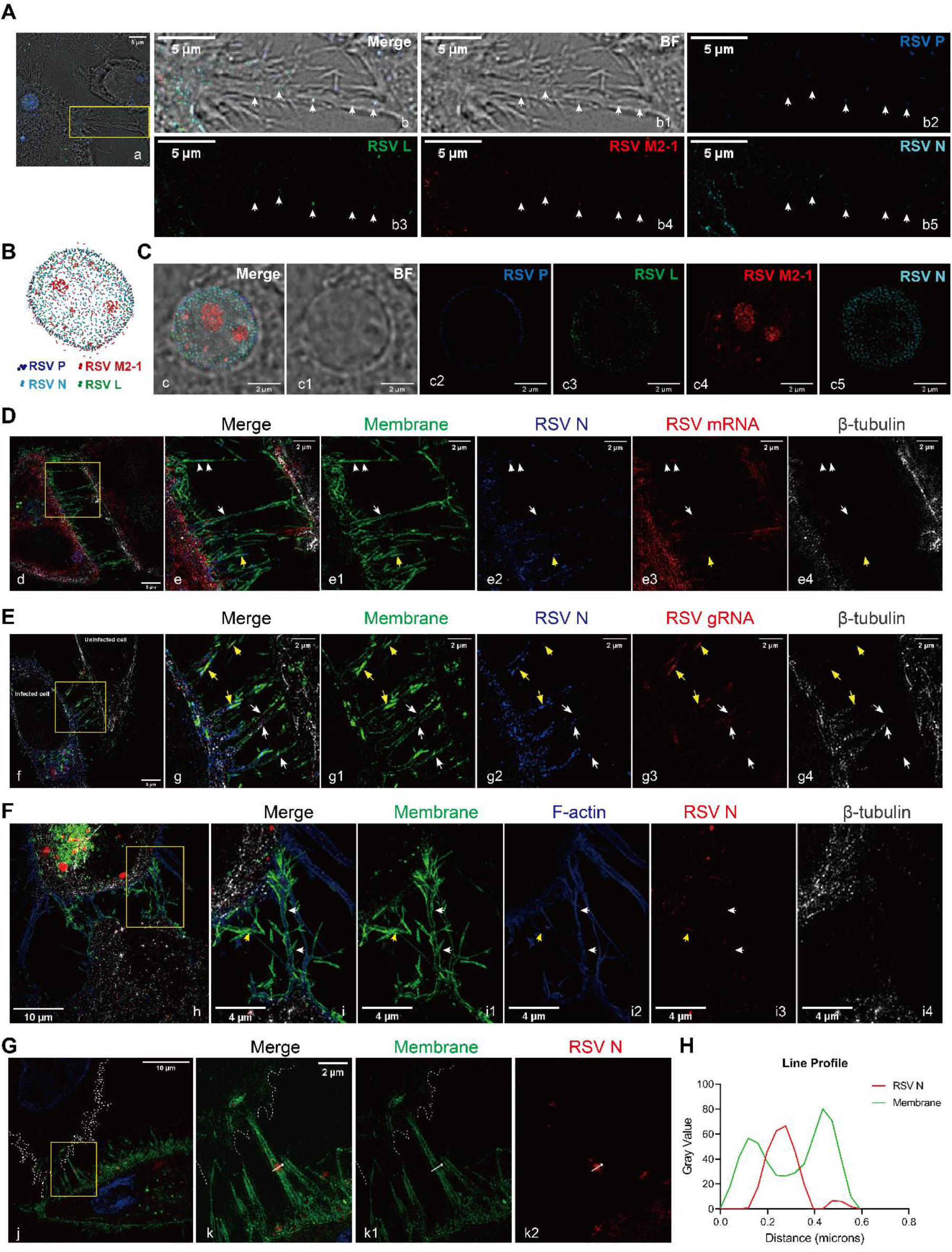
Compositions of the intercellular connected nanotubes and viral materials. Infected cells were fixed and then stained with various antibodies targeting viral proteins or probes targeting viral RNAs before being imaged in ultrahigh-resolution mode. (A) Composition of viral materials located in intercellular connected nanotubes. (B-C) Composition of IBs. (D-E) Viral genome RNAs and mRNAs located in intercellular connected nanotubes. (F) The cytoskeleton participates in the formation of connected nanotubes. (G) Intercellular connected nanotubes might be derived from infected cells. (j-k1) Infected cells and tunnels had strong signals of WGA binding, but uninfected cells did not (the membrane is shown by the white dotted line). (k2) IBs in intercellular connected nanotubes. (H) Line profile of an IB in (G). The white notched arrows indicate viral materials located in the tunnels, and the yellow filled arrows indicate virions on the surfaces of filaments. The following source data is available for figure 4: **Source data 1.** This spreadsheet contains data used to generate graphs in Figure 4H.

To capture clearer spatiotemporal evidence of the possible spread mode, the IBs were marked by infecting the cells before transfection with a GFP-fused N gene. With live cell imaging, we found that IBs were dynamic; they continually merged and split (Figure 5A). Under the ultrahigh-resolution mode, we observed the direct transfer of IBs through intercellular connected nanotubes in the early stage of infection. IBs from infected cells were rapidly transferred to neighboring cells, and the whole event took only approximately 4 min (Figure 5B).

**Figure 5.**
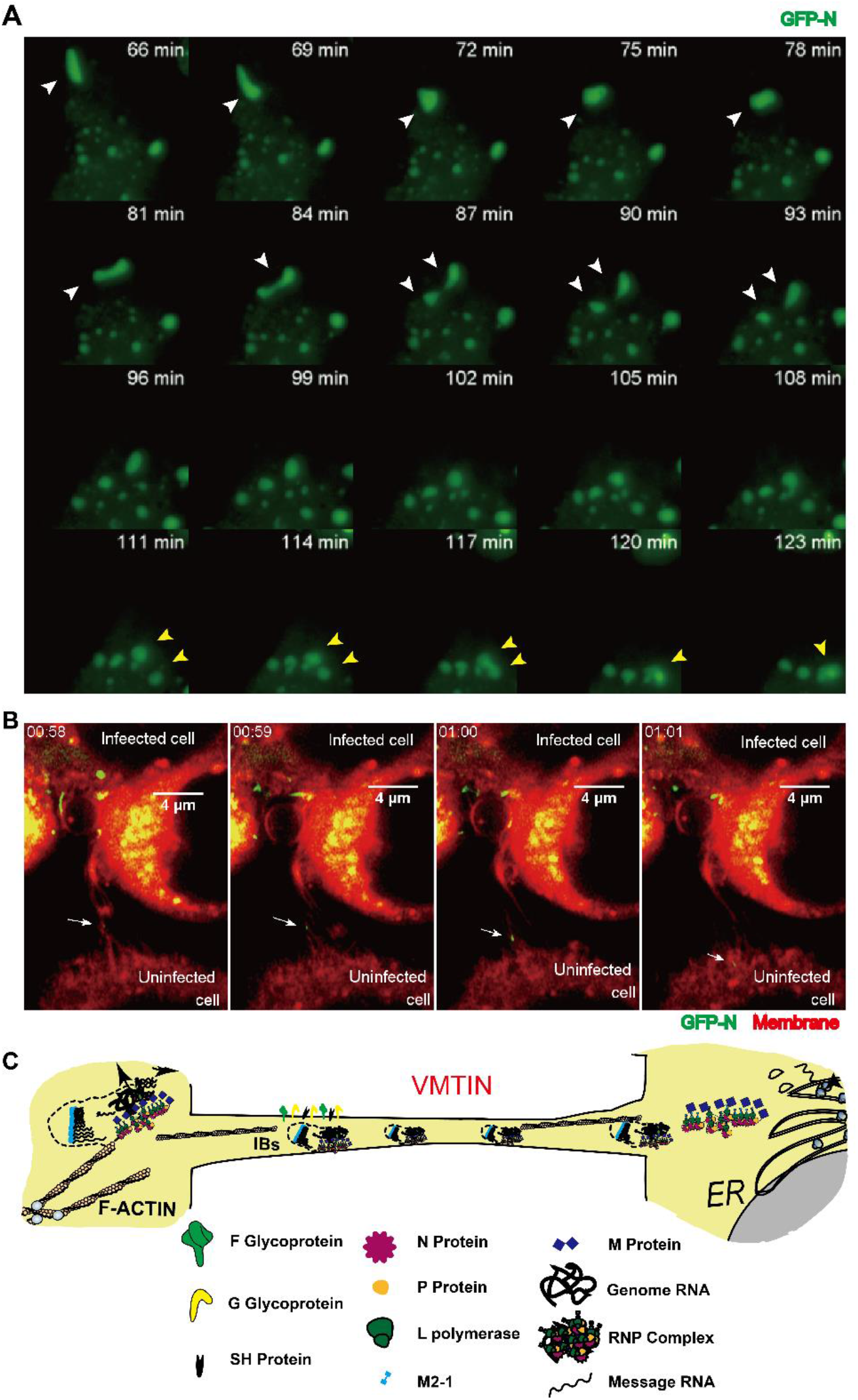
Direct migration of viral materials (IBs) from cell to cell through intercellular connected nanotubes. IBs were marked by infecting the cells before transfection with a GFP-fused N gene, and monitoring of IB movement was completed in the early stage of infection (within 11 h.p.i.). (A) Merging and fission of IBs. The white arrows show the fission of IBs, the yellow arrows show the merging of IBs, and the GFP-N signal (green) represents the IBs. (B) Migration of IBs from infected cells to uninfected cells. The white arrow represents a shifting IB, the DID signal (red) represents the membrane, and the green signal represents IBs. (C) Sketch of the VMTIN mode.

Based on the evidences obtained thus far, a model of the intricate mechanism of RSV material diffusion was created (Figure 5C). First, infected cells might protrude filopodia to neighboring cells that become open-ended intercellular nanotubes, in which microfilaments participate. Second, viral materials, probably in the form of IBs (replication factories), are transferred from infected cells to neighboring cells prior to virion budding (from 3 h.p.i.) and throughout the whole infection period. Third, virion-based spread, including cell-free virion dissemination and filopodia-driven virion intercellular delivery, starts to occur after virion assembly (from 11 h.p.i.). We named the new spread mode “viral material transfer through intercellular nanotubes” (VMTIN).

### Direct migration of viral materials induced effective infection despite the presence of high-titer antibodies

How does the viral material transfer through TNT-like nanotubes affect RSV infection? A key question to answer was whether the transfer of viral materials into the target cells successfully induced replication and expression. To answer this question, we utilized another coculture system like that described above; the only difference was in the detection step. At six hours post mixing, mKate-DIO+ cells were sorted and further cultured for another twenty-four or forty-eight hours (Figure 6A). According to the results of statistical analysis, the number of mKate+DIO+ cells significantly increased, and RSV genomic RNA was amplified in these cells (Figure 6B-6D). In summary, the migration of viral materials to receptor cells successfully induced infection, which could not be completely inhibited by neutralizing antibodies.

**Figure 6.**
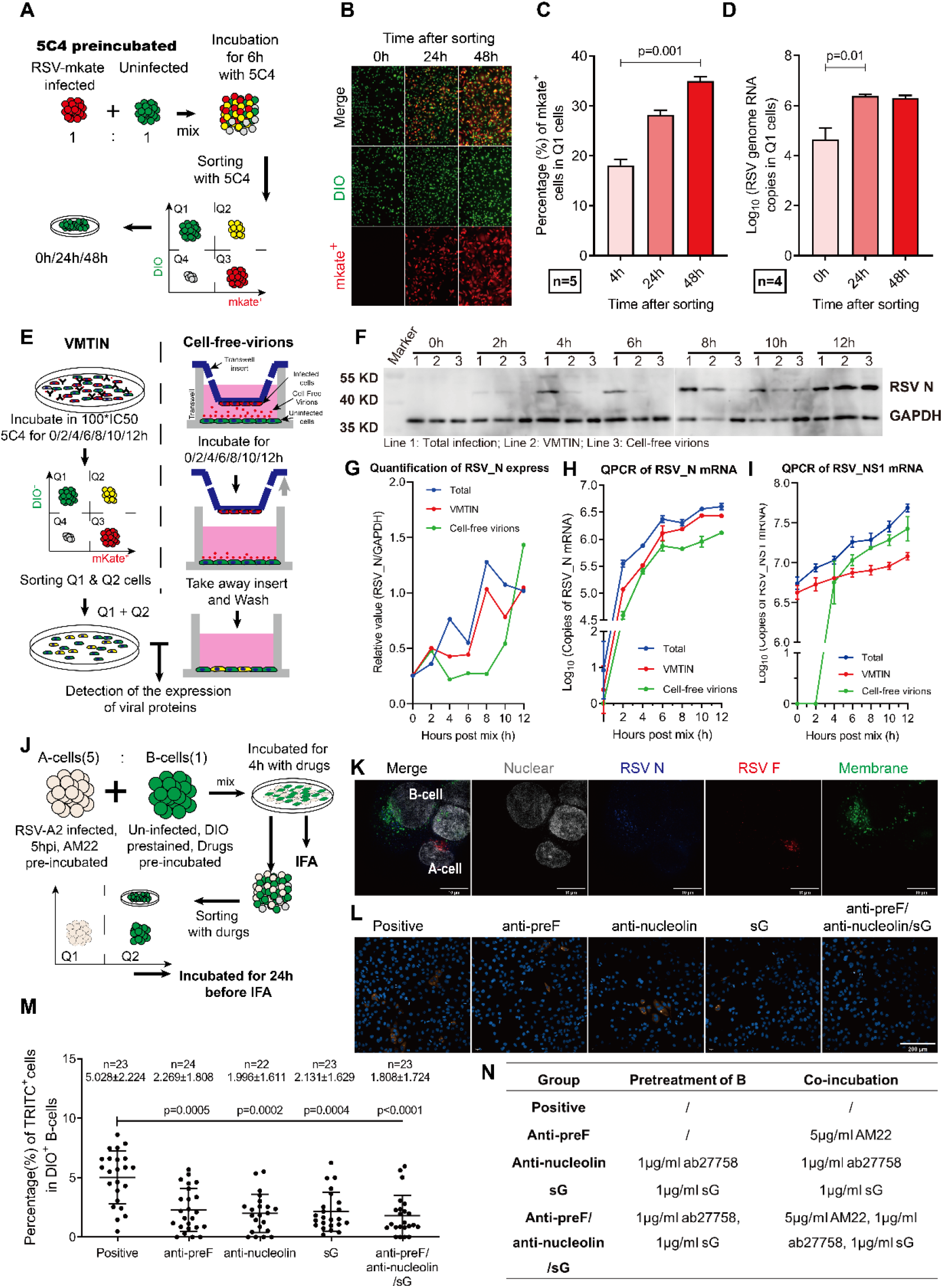
VMTIN successfully and efficiently induced infection. (A-D) VMTIN induced RSV replication in receptor cells. RSV-A2-mKate-infected cells (donor cells, mKate+) and uninfected cells (receptor cells, DIO^+^) pretreated with 100 IC50 5C4 were mixed and further incubated. Subsequently, DIO^+^mKate^-^ cells were sorted for further culture and analysis. (A) Sketch of the experimental scheme. (B) Fluorescence microscopic images of sorted DIO^+^mKate^-^ cells. (C) Quantitative analysis of images in (B). (D) Detection of RSV genomic RNA in sorted cells. (E-H). VMTIN is highly efficient. RSV-A2-mKate-infected cells were preincubated with 100 IC50 5C4 and digested. For VMTIN, the infected cells (donor cells, mKate^+^) were then mixed and further cultured with uninfected cells (receptor cells, DIO^+^) in the presence of 5C4. For cell-free virions, the infected cells were seeded on the back of the membrane of a Transwell insert, while uninfected cells were seeded in the bottom of the Transwell chamber. After further culture, samples (DIO^+^ and DIO^+^mKate^+^ cells) were collected for further analysis. (E) Sketch of the experiment. (F) Western blot of samples. (G) Quantitation of (F). (H-I) QPCR of samples. (J-N) VMTIN was partly inhibited by blocking the function of F or G. RSV-A2-infected cells were pretreated with 100 IC50 AM22 and named A-cells. Uninfected cells were pretreated with various drugs and then prestained with the dye DIO; these cells were named B-cells. A-cells and B-cells were mixed and further cultured in the presence of various drugs. Four hours later, parts of the mixtures were examined for virions adherent to cell membranes via immunofluorescence (IF), and the rest of the mixtures were sorted to collect DIO^+^ cells, which were used to determine the infection rate after further culture. (J) Sketch of the experiment. (K) IF to ensure the absence of virions on cells. (L) IF to determine the infection rate. (M) Quantitation of (K). (N) Drugs and their doses. The results are presented as the geometric mean ±SD. The following source data is available for figure 6: **Source data 1.** This spreadsheet contains data used to generate graphs in Figure 6C-D, Figure 6G-I, 6M.

We next sought to evaluate the relative contribution of VMTIN to RSV spread in monolayer culture. In the coculture group, both mKate-DIO+ cells and mKate+DIO+ cells were collected after several hours of incubation. In the control group, a Transwell system was utilized to mimic cell-free virion dissemination. We first seeded donor cells on the lower surface of the Transwell insert and then inserted the insert into a Transwell chamber that was preseeded with receptor cells (Figure 6E). After a period of incubation, the insert was removed, and the receptor cells in the bottom of the chamber were washed three times and collected for further detection. From the results (Figure 6F-6H), we found that the start of N protein transcription and expression induced by VMTIN in the presence of 5C4 occurred more quickly than that of cell-free virions but was slightly slower than total spread.

As mentioned above, VMTIN was not completely inhibited by the pre-F-specific neutralizing antibody, but whether viral proteins indeed participated in this process remained unclear. We examined the influences of several drugs on VMTIN (Figure 6J-6N), including AM22 (another high-titer neutralizing antibody that specifically targets pre-F), ab27758 (an antibody targeting the receptor of F, nucleolin) and sG (the secreted form of G protein). First, A-cells were infected with RSV-A2 for 5 h and pretreated with AM22 at a concentration of 5 μg/ml, while B-cells were prestained with the dye DIO and pretreated with different drugs. Thereafter, A-cells and B-cells were digested and resuspended in drug-containing medium and then mixed at a ratio of 5 to 1. The mixture was incubated for another 4 h. After that, part of the mixture was examined for the existence of virions by immunofluorescence (IF) (Figure 6K), and then DIO+ cells were sorted from the rest of the mixture and further incubated for another 24 h. Finally, the sorted DIO+ cells were examined, and the newly infected cells were counted. The results showed that these three drugs suppressed VMTIN, while their synergistic effect was not significant (Figure 6L-6M). The results suggested that the F protein and G protein might play roles in VMTIN.

### DVGs probably utilize VMTIN to disseminate among cells, which might contribute to the persistence of latent infection

RNA virus replication is error-prone, and DVGs might accumulate during the process (Ziegler and Botten, 2020). Therefore, some infected cells gradually accumulate high levels of DVGs but have low levels of full-length genomes (FLGs) (Figure 7—figure supplement 1). These cells are known as DVG-high cells and might establish latent persistence of RSV (Xu et al., 2017, Valdovinos and Gómez, 2003). We were interested in the connections between the DVG-high cells and neighboring cells and whether these connections would aid latent persistence. First, we produced high-DVG RSV virions via infection with a high MOI. Then, we compared the characteristics of low-DVG RSV-infected (DVG-low) cells and high-DVG RSV-infected (DVG-high) cells by IF-fluorescence in situ hybridization (FISH). According to the data, DVG-low cells expressed high levels of viral proteins and produced numerous filamentous virions via budding from the plasma membrane, while DVG-high cells expressed extremely low levels of viral proteins and produced scarce virions despite the presence of IB-like structures (Figure 7A). We also found TNT-like intercellular nanotubes between DVG-high cells and neighboring cells, in which viral materials were located (Figure 7B).

**Figure 7.**
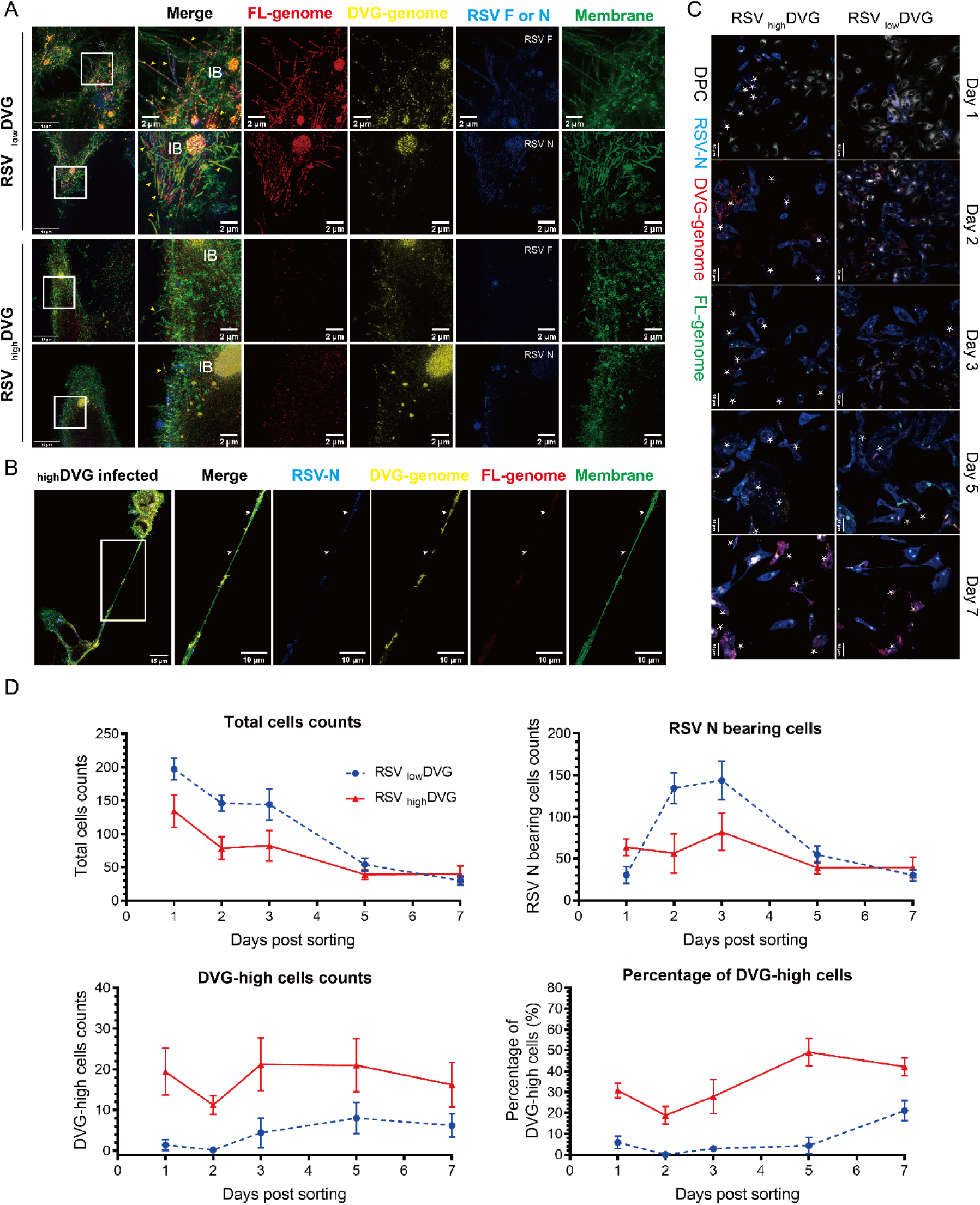
High-DVG RSV utilized VMTIN but not virion-based modes to spread among cells, which might contribute to the persistence of latent infection. (A-B) Cells infected with high-DVG RSV or low-DVG RSV were examined for the expression of viral proteins and viral RNA via IF-FISH. (A) Comparison of features between DVG-high cells and DVG-low cells; the yellow arrows indicate progeny virions, and the scale bar represents 2 μm. (B) Viral materials in the intercellular nanotubes between DVG-high cells and neighboring cells. The white arrows indicate the viral material location signals; the scale bar represents 10 μm. (C-D) High-DVG RSV might contribute to the persistence of latent infection. Cells infected with high-DVG RSV or low-DVG RSV were digested at 5 h.p.i. and then mixed with uninfected cells (prestained with DIO). After 4 h of coculture, DIO^+^ cells were sorted from the mixture and further cultured. Some DIO^+^ cells were fixed and examined for viral protein expression as well as viral RNAs in the following days. (C) Detection of viral proteins and viral RNAs in sorted DIO+ cells by IF-FISH. The white stars indicate DVG-high cells; the scale bar represents 50 μm. (D) Quantitation of specific cells among sorted cells. The following source data is available for figure 7: **Source data 1.** This spreadsheet contains data used to generate graphs in Figure 7D. **Figure supplement 1.** DVG-high cells gradually appeared in routine infection with low-DVG RSV.

The lack of virion formation on the cell membrane suggested that cell-free virion dissemination and virion transfer to adjacent cells hardly occurred. Thus, we were interested in whether and how DVG-high cells could induce infection of neighboring cells. We mixed low-DVG RSV or high-DVG RSV (MOI=4, 5 h.p.i.)-infected donor cells with uninfected receptor cells prestained with the membrane dye DIO. After 4 h of coculture, DIO+ cells were sorted and incubated without neutralizing antibodies. As shown in Figure 7C-7D, after 24 h of incubation, some sorted DIO+ cells began expressing the RSV N protein in both groups, implying that DVG-high cells could spread in the absence of virions. The number of RSV-infected cells increased drastically in the low-DVG RSV group but only slightly in the high-DVG RSV group (Figure 7D), implying the weak infection ability of high-DVG RSV. The cells in the low-DVG RSV group were mainly DVG-low in the first several days, while DVG-high cells gradually appeared in the following days, although very few appeared. Some cells in the high-DVG RSV group were DVG-high, the number of which remained basically the same. Cells in both groups gradually died after sorting, especially the DVG-low cells, which seemed to die more easily than the DVG-high cells. Overall, the results indicate that DVG-high cells probably induce infection in neighboring cells in the absence of virions via VMTIN, which might promote the establishment of the latent persistence of RSV, at least in monolayer cells.

## Discussion

Neutralizing antibodies are key to the protection provided by many vaccines, and RSV vaccines are no exception. One of the reasons for the ERD caused by FI-RSV was thought to be the lack of pre-F in the vaccine (Killikelly et al., 2016) and the subsequent inability to induce high titers of neutralizing antibodies (Murphy et al., 1986, Murphy and Walsh, 1988, Ngwuta et al., 2015). However, in a recent report, a pre-F subunit vaccine that did induce pre-F-specific high-titer neutralizing antibodies still caused ERD at low doses (Schneider-Ohrum et al., 2017, Phung et al., 2019). In addition, it was noteworthy that when the serum neutralization level dropped to a certain critical point, breakthrough infections began to occur and then become out of control, eventually leading to severe lung pathology in the vaccine-immunized animals despite the immunological memory induced by the vaccine. This raised the question of whether the virus can bypass the immune system and spread, preventing it from being contained in the event of breakthrough infection in vaccinated animals. To answer this question, we investigated the intercellular transmission of RSV.

Paramyxoviruses can spread from cell to cell using a variety of different mechanisms (Cifuentes-Muñoz and Ellis Dutch, 2019). The syncytium is formed by cell fusion induced by a fusion protein (Shigeta et al., 1968), and the mature progeny virions are released from the cell body; this is the classic cell-to-cell transmission mode of paramyxoviruses. In recent years, some new modes of transmission have also been proposed; for example, the measles virus is believed to be able to transfer materials between cells by forming fusion pores through local fusion (Singh et al., 2015), and human metapneumovirus (HMPV) is speculated to be able to transfer viral proteins through intercellular extensions (mainly based on the localization signals of viral proteins observed in intercellular extensions) (El Najjar et al., 2016). The understanding of RSV transmission was previously limited to the classical modes. However, for RSV, large numbers of syncytia could not be observed in tissue (Neilson and Yunis, 1990), and most of the progeny virions produced were not spontaneously released from the cell membrane (Collins et al., 2013b). Only approximately 5% of the virions were released into the extracellular space as free virions. In addition, due to the fragility of virions or the randomness of their movements, the propagation efficiency of cell-free virions was low. In a recent study, it was found that intercellular filopodia directly delivered virions that had not been released from RSV-infected cells to neighboring cells (Mehedi et al., 2016), and this multicopy delivery was certainly efficient.

However, the aforementioned RSV transmission modes depended on the F protein or virions and could theoretically be blocked by neutralizing antibodies. Interestingly, we found that intercellular spread of RSV still occurred in the presence of high concentrations of neutralizing antibodies after blockade of cell-free virion and filopodia-driven intercellular virion delivery (Figure 1). In addition, we found that viral proteins were located in the open-ended intercellular nanotubes between infected and uninfected cells, which pointed to a possible new mechanism of intercellular diffusion in which infected cells transferred viral materials to neighboring cells via intercellular nanotubes. We further observed this transfer process directly through ultrahigh-resolution live cell imaging. We also found that this transmission occurred in the early stages of viral infection, before the formation of virions, and was the major transmission event in the early stages of infection. The early onset and neutralizing antibody insensitivity of the new mode of transmission increased its efficiency, which might have been part of the reason why the large numbers of antibodies produced by the activation of immune memory might have been unable to control the spread of the virus once breakthrough infection occurred.

The viral material transmitted by this mode might not be selective, and it was determined to include the components of IBs (N protein, P protein, L protein, M2-1 protein, genomic RNA, and mRNA) as well as the membrane fusion protein (F protein). IBs are considered the replication factories of RSV (Rincheval et al., 2017), and the transfer of IBs should be highly efficient in theory, as they have a great advantage over cell-free virions. We demonstrated that in several RSV transmission events, the new mode of transmission initiated viral protein expression more efficiently than cell-free virion diffusion. The apical filopodia of virus-infected cells are usually distributed with multiple virions, so filopodia-driven virion delivery is usually multicopy; the HIV-1-related hypothesis (Boullé et al., 2016) thus implies that this mode of transmission should have high efficiency, but it was not the same as the efficiency of the newly discovered mode of transmission in this study.

Viral infection usually hijacks the cytoskeleton for the viral life cycle. Viral infection might increase cell motility, which is generally thought to be due to viral regulation of the cytoskeleton, but this feature is not evident in HMPV and parainfluenza virus type 3 (PIV3) (Mehedi et al., 2016). Arp2 was involved in the motility enhancement of RSV-infected A549 cells, which was also demonstrated in RSV-infected HEp-2 cells by our wound healing experiments and motility assays. This motility enhancement might have promoted the cell-to-cell contact frequency, which probably increased the transfer of virions. Viral infection can also cause rearrangement of the cytoskeleton, which changes cell morphology. HMPV infection promotes increased generation of intercellular extensions (El Najjar et al., 2016). and RSV infection can induce host cells to produce more filopodia through ARP2 regulation (Mehedi et al., 2016). In this study, we found that in addition to increasing filopodia formation, RSV infection promoted the generation of open-ended intercellular nanotubes, which were probably derived from filopodia of infected cells (the glycoprotein staining characteristics of nanotubes were the same as those of infected cells but not uninfected cells). The skeletons of these nanotubes were mainly composed of F-actin rather than β-tubulin, similar to the TNTs between HIV-infected immune cells (Eugenin et al., 2009).

DVGs are produced during the replication of most RNA viruses (Lazzarini et al., 1981, Ziegler and Botten, 2020). DVGs have been shown to promote the survival of infected cells through a MAVS/TNFR2-mediated mechanism (Xu et al., 2017, Dakhama et al., 1997). In addition, DVGs can stimulate the innate immune response and interfere with viral genome replication (Ziegler and Botten, 2020, Sun et al., 2015), thus reducing the severity of disease. We found that cells containing high levels of DVGs (DVG-high) expressed reduced levels of viral antigens and produced almost no progeny virions, suggesting that DVG-high cells might not spread through the classical virion-dependent mode. Viral proteins were also observed to be localized in intercellular nanotubes by ultrahigh-resolution imaging, suggesting that DVG-high cells may also conduct cell-to-cell diffusion through this mode, which might contribute to the establishment of persistent latent RSV infection.

## Acknowledgements

We thank Dr. Peter Franklin and Liu Qing-Feng for the support on ultrahigh-resolution imaging. We thank Dr. Barney S. Graham (VRC, NIH) for the generous donation of the recombinant virus RSV-mKate (pSynkRSV-A2-D46F). This research was funded by National Natural Science Foundation (Grant number 31730029, 81991491, 82071783) and CAMS Innovation Fund for Medical Sciences (Grant number 2019RU022).

## Author Contributions

Conceptualization, Z.W., Z.Z.Z. and X.N.S.; Methodology, Z.W. and L.X.; Validation, L.X., Z.L.J, and S.J.Y.; Formal Analysis, Z.W. and L.X.; Investigation, L.X., Z.W., Z.L.J., C.L. and W.G.H.; Writing-Original Draft, Z.W. and L.X.; Writing-Review & Editing, Z.W. and Z.Z.Z.; Funding Acquisition, X.N.S. and Z.Z.Z.; Resources, L.X., S.J.Y., Z.M., C.L., S.Y.P., Z.W. and Z.L.T.; Supervision, W.Y.B. and Z.Z.Z.

## Materials and Methods

### Cells and viruses

HEp-2 cells (ATCC, USA) were cultured with Minimum Essential Medium (Gibco, USA), which was supplemented with 10% (v/v) fetal bovine serum (FBS) (Gibco). RSV strain A2 (ATCC) and RSV-A2-mKate (a recombinant virus expressing a fluorescent mKate protein) were propagated in HEp-2 cells using MEM supplemented with 10% (v/v) FBS. The virus titers were measured by plaque assay.

### Plasmids, antibodies and reagents

To label ribonucleoprotein (RNP) complexes or IBs, a plasmid encoding the EGFP-N fusion protein was constructed and transfected into infected HEp-2 cells with Lipo3000 (Thermo Fisher, USA) according to the manufacturer’s instructions. Various antibodies were used in this study, including 11H8 (anti-RSV N, a mouse ascites antibody), 15A3 (anti-RSV P, a mouse ascites antibody), 14A10 (anti-RSV L, a mouse ascites antibody) and RSV 5H5 (anti-M2-1, ab94805) (Abcam, UK). All primary antibodies were used at 10 μg/ml. Alexa Fluor 647-conjugated EPR16774 (anti-β-Tubulin, ab204686) (Abcam) and fluorescein isothiocyanate labeled phalloidin (Sigma-Aldrich, USA) were used to label the cytoskeleton at the recommended concentrations. The secondary antibodies used were donkey anti-mouse Alexa Fluor 405 (Abcam), donkey anti-mouse TRITC (Sigma-Aldrich), goat anti-human Alexa Fluor 546 (Thermo Fisher), and goat anti-rabbit Alexa Fluor 405 (Thermo Fisher). All secondary antibodies were used at 5 μg/ml.

For colocalization of viral proteins, 11H8, 14A10 and RSV 5H5 were labeled using a Zenon Tricolor Mouse IgG1 Labeling Kit according to the manufacturer’s protocol, and the reagents were used at 10 μg/ml. 15A3 was labeled using EZ-Link Sulfo-NHS-LC-Biotin (Pierce, USA) according to the manufacturer’s protocol, and the reagent was used at 5 μg/ml. Streptavidin allophycocyanin conjugate (Life Technologies, USA) was used at 2 μg/ml. Wheat Germ Alexa Fluor 488 (Invitrogen, USA) was used at 10 μg/ml.

For Western blotting, the primary antibody (11H8) used was diluted to 1 μg/ml, and anti-GAPDH (rabbit polyclonal) (Proteintech, USA) was diluted to 0.2 μg/ml. The secondary antibodies were goat anti-rabbit HRP (Invitrogen) and goat anti-mouse HRP (Abcam, UK) diluted to 0.2 μg/ml.

### IF

HEp-2 cells were infected with RSV-A2 (MOI=4). Twenty-four hours later, the digested infected cells were mixed with uninfected cells at a ratio of 1:3, and then the mixtures were seeded onto glass coverslips (2.5×105 cells per well) in six-well plates for further incubation. Eighteen hours later, the cells were fixed with PBS-formaldehyde 4% (w/v) for 15 min at 37 °C, washed with PBS and permeabilized with PBS-Triton X-100 0.2% (v/v) for 10 min at room temperature (RT). The cells were blocked for 1 h in PBS-skim milk 5% (w/v) at 4 °C, washed with PBS and incubated with primary antibodies at RT for 1 h. Then, the cells were incubated with secondary antibodies at RT for 1 h. The nuclei were stained with DAPI (Invitrogen). Coverslips were then mounted with SlowFade Gold Antifade Mountant (Invitrogen).

### In situ hybridization

Cells were fixed with neutral buffered formalin (NBF) 10% (v/v) for 30 min at RT. Then, pretreatment and FISH were performed using an RNAscope Multiplex Fluorescent Reagent Kit v2 (ACDBio, USA) following the manufacturer’s instructions. Briefly, samples were pretreated with hydrogen peroxide for 10 min at RT, washed with PBS, and treated with protease III (1:15 dilution in PBS) for 10 min at RT. The samples were hybridized with probes that targeted specific sequence positions of RSV genomic RNA and mRNA for 2 h at 40 °C in a HybEZ hybrid system (ACDBio). RNAscope detection reagents were used to amplify the hybridization signals via sequential hybridization of amplifiers. Opal 570 Reagent (PerkinElmer, USA) and Opal 670 Reagent (PerkinElmer) were used to stain the different channels at dilutions of 1:1500 in Multiplex TSA Buffer (ACDBio). Coverslips were then mounted with SlowFade Gold Antifade Mountant (Invitrogen). The sequences of the probes used for in situ hybridization were the same as those in a recent report (Xu et al., 2017).

### Ultrahigh-resolution microscopy

IF and in situ hybridization images were acquired on a DeltaVision OMX V4 (GE Healthcare, USA) equipped with a 60×/1.42 NA PlanApo oil immersion objective (Olympus, Japan). Images were acquired with OMX Master (GE Healthcare). Image registration was performed in softWoRx 6.1.1. The voxel size of the reconstructed images was 0.125 μm in the z-axis.

### Wound-healing assay

HEp-2 cells were plated in six-well plates (5× 105 cells per well) and grown overnight at 37 °C. Then, the cells were mock-infected or infected with RSV-A2 at an MOI of 4. Twenty-four hours later, the cell monolayers were scratched with a 200 μl pipette tip and then imaged every 20 min for 12 h with Harmony High-Content Imaging and Analysis Software version 4.9 (PerkinElmer) and an Opera Phenix High Content Screening System (PerkinElmer) with a 5×/0.16 NA objective in an environmental chamber with CO2 at 37 °C. Six randomly selected locations were imaged per sample. Cell migration was measured by quantifying the number of mKate+ cells in the wound and determining the area of the wound using Fiji-ImageJ (National Institutes of Health, USA). Cell migration tracking was analyzed with the Columbus Image Data Storage and Analysis System.

### Antibody inhibition assays for different modes of RSV spread among cells

Antibody inhibition assay for cell-free-virion infection. HEp-2 cells (5× 105 cells per well) in six-well plates were infected with RSV-A2-mKate at an MOI of 4. At 24 h.p.i., the cells were washed with PBS. Then, the cells were incubated at 37 °C in 2 ml of complete medium with 0 μg/ml, 0.1 μg/ml, 1 μg/ml, 5 μg/ml, or 10 μg/ml 5C4 (anti-RSV F, mouse monoclonal). At 6 h, 12 h, and 48 h post 5C4 addition, the supernatant of the tumor cell culture was harvested and centrifuged at 162 ×g for 3 min. After centrifugation, 3×104 HEp-2 cells per well in 96-well plates were incubated with 200 μl of supernatant for 24 h. Fluorescence imaging was performed with a Nikon Eclipse Ti-S inverted microscope epifluorescence microscope equipped with a 10× CFI60 objective (Nikon, Japan) and a DS-Ri1 camera. Antibody-mediated inhibition of cell-free virion infection was measured by calculating the number of mKate+ cells in randomly selected locations imaged per well.

Antibody inhibition assay for virion-based transmission. HEp-2 cells (5×105 cells per well) in six-well plates were infected with RSV-A2 at an MOI of 4. At 24 h.p.i., the culture medium was removed, and 2 ml of MEM with 0 μg/ml or 1 μg/ml 5C4 was added. After incubation and PBS washing, the cells were scraped in 1 ml of MEM with 0 μg/ml or 1 μg/ml 5C4. Cells that had been plated onto coverslips at a density of 2.5× 105 cells per well in six-well plates and grown overnight at 37 °C were incubated with 0.5 ml of liquid containing the scraped cells for 1 hour on a shaking table at RT. After washing with PBS 3 times, the cells on coverslips were incubated with 2 ml of complete medium with 0 μg/ml or 1 μg/ml 5C4. At 2 h and 16 h after the liquid containing the scraped cells was added, the cells were fixed, stained and imaged.

Coculture assay for direct migration of viral materials from cell to cell. HEp-2 cells (5×105 cells per well) in six-well plates were infected with RSV-A2-mKate (MOI=4). At 24 h.p.i., the culture medium was removed, and 2 ml of MEM with 1 μg/ml 5C4 was added. After incubation and PBS washing, the cells were digested with MEM with 1 μg/ml 5C4. A total of 5×105 naïve HEp-2 cells were stained with Vybran DIO Cell-Labeling Solution (0.005 μg/ml, Thermo fisher) for 20 min at 37 °C and washed with PBS. Then, the infected cells were mixed with naïve cells stained with DIO in a 1:1 ratio in MEM with 1 μg/ml 5C4 (every 5×105 infected cells needed 2 μg of 5C4). The cells were cultured, and the culture medium was changed every 6 h.

### Flow cytometry analysis

Cells were washed with PBS, trypsinized with 0.25% trypsin-EDTA (Gibco), incubated with culture medium to neutralize trypsin activity, and centrifuged at 162 ×g for 3 min. The cells were washed and resuspended in PBS. The cell suspension was passed through a 70-μm cell strainer, and the cells were subjected to analysis and sorting using a FACSAria III flow cytometer (BD Biosciences, USA). HEp-2 cells infected with RSV-A2-mKate at an MOI of 4 and naïve DIO-stained cells were used for compensation, and a fluorescence minus one control was included to aid in setting gates. Acquisition was performed until 50,000 cells were recorded. Gating was performed on live singlet cells followed by DIO+mKate+ cells using FACSDiva software (BD Biosciences). Analysis was performed with FlowJo software version 10.5.0 (BD Biosciences).

### Western blot analysis

Cells were centrifuged at 162 ×g for 3 min. The sediments were lysed with Cell Lysis Buffer for Western and IP (Beyotime, China) with 1 mM phenylmethanesulfonyl fluoride (Beyotime); every 106 cells needed 200 μl of lysis buffer. Then, the samples were centrifuged at 17000 ×g for 5 min. Then, 100 μl of supernatant was mixed with 6× SDS loading buffer, boiled for 15 min at 100 °C and loaded into the wells of a gel alongside a molecular weight marker (Thermo fisher). The gel was run in a Mini-PROTEAN Tetra Cell system (Bio-Rad, USA) in 1× SDS running buffer at a constant voltage of 80 V for 120 min. Then, the proteins were transferred to 0.45 μm pore nitrocellulose filter membranes (Whatman, UK) in Towbin system buffer (25 mM Tris-HCl pH 8.3, 192 mM glycine, 20% (v/v) methanol) at a constant amperage of 300 mA for 90 min using a Mini Trans-Blot Cell (Bio-Rad). The membranes were incubated in NRA blocking buffer (INNOVAX, China) for 1 hour at RT and then with primary antibodies in ED-13 buffer (INNOVAX). The membranes were washed 5 times for 5 min each in PBST (PBS with 0.2% Tween-20) and then incubated with secondary antibodies for 1 hour. After washing, the membranes were imaged using an ImageQuant LAS4000 mini (GE Healthcare).

### Preparation of low-DVG virus and high-DVG virus

Viruses containing different levels of DVGs were produced according to methods in recent studies (Sun et al., 2015, Xu et al., 2017).

### Statistical analysis

The results were plotted and statistical analysis was performed using GraphPad Prism 9.00 (GraphPad, USA). Two populations data was performed with Mann-Whitney test. Three or more populations data (one independent variable) was performed Kruskal-Wallis test and Dunn’s multiple comparisons test. Three or more populations data (two independent variables) was performed Oridinary two-way ANOVA and Šídák’s multiple comparisons test. Only a p value <0.05 was considered to indicate statistical significance. Confidence interval was 95%.

## Supplemental information

**Figure 2—figure supplement 1.**
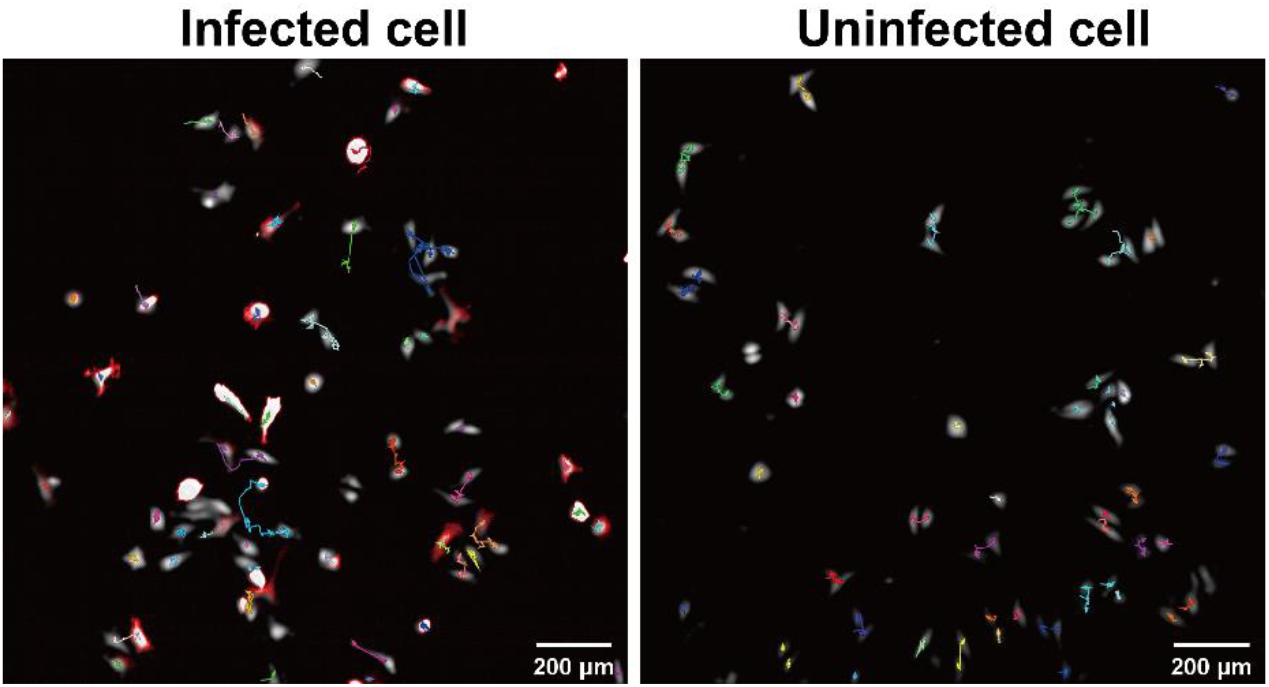
RSV infection increased cell movement. RSV-A2-mKate infected cells (red) were tracked for the movement with Opera Phenix High Content Screening System, the migration paths were showed.

**Figure 2—figure supplement 2.**
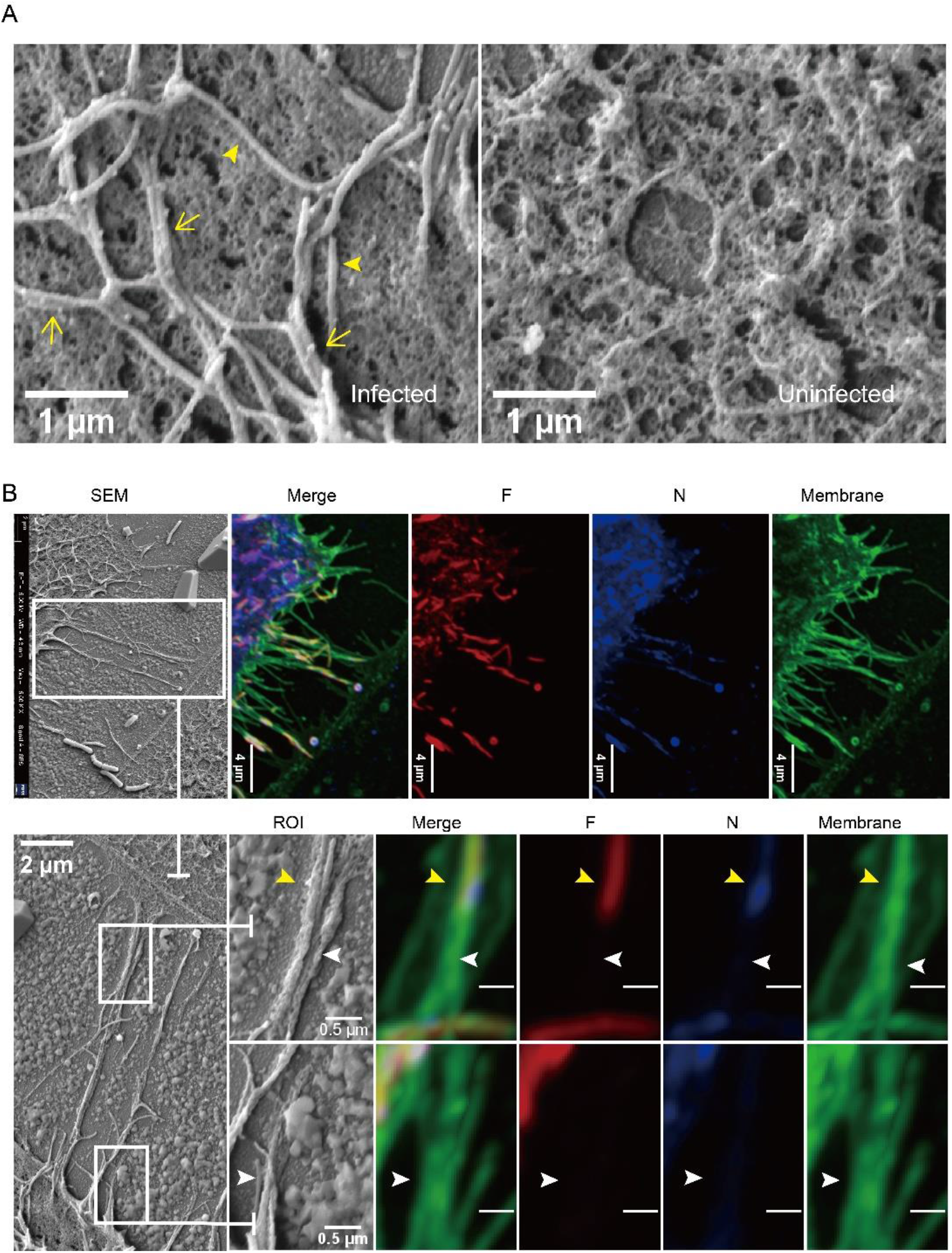
The morphological features of infected cells under the CLEM. RSV-A2 infected cells were treated with CLEM system. (A) The comparison of the cell surface of infected cell and uninfected cell under scan electron microscopy. Yellow filled arrows represent the closed virions, and yellow opened arrows represent the opened virions. (B) The morphology of intercellular connected nanotubes and the related viral materials distribution under CLEM.

**Figure 2—figure supplement 3.**
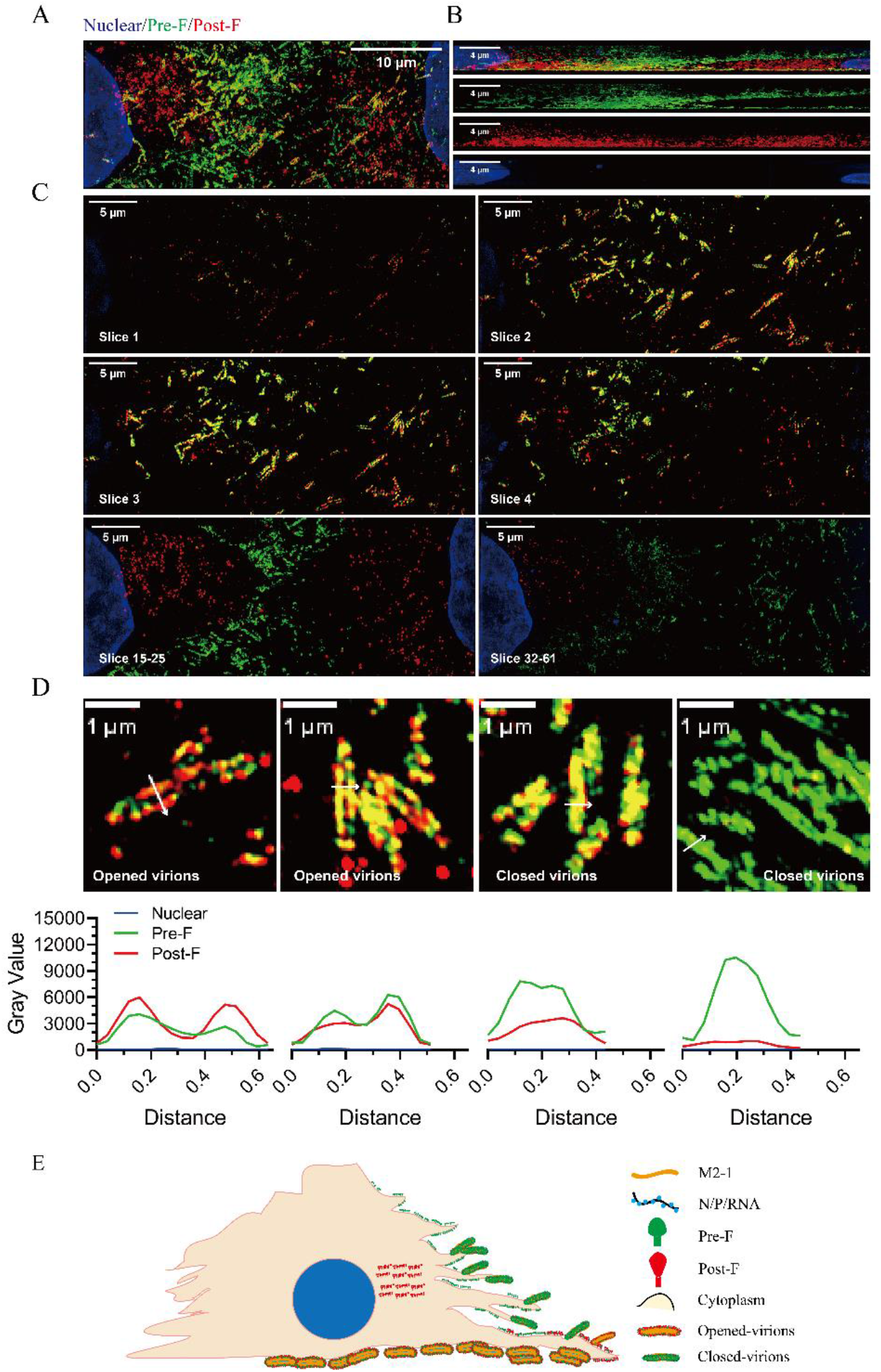
The distribution and the abundance of pre-F and post-F on infected cells. RSV-A2 infected cells were co-stained with antibodies specially targeting pre-F or post-F, and the images were obtained with a ultrahigh resolution microscopy. (A) The overview of the infected cells, which was obtained via executive the Z-projection of 61 slices. (B) The side view of (A). (C) Several chose slices showing distribution of pre-F and post-F. (D) Various virions containing different level pre-F. (E) The sketch of infected cells.

**Figure 7—figure supplement 1.**
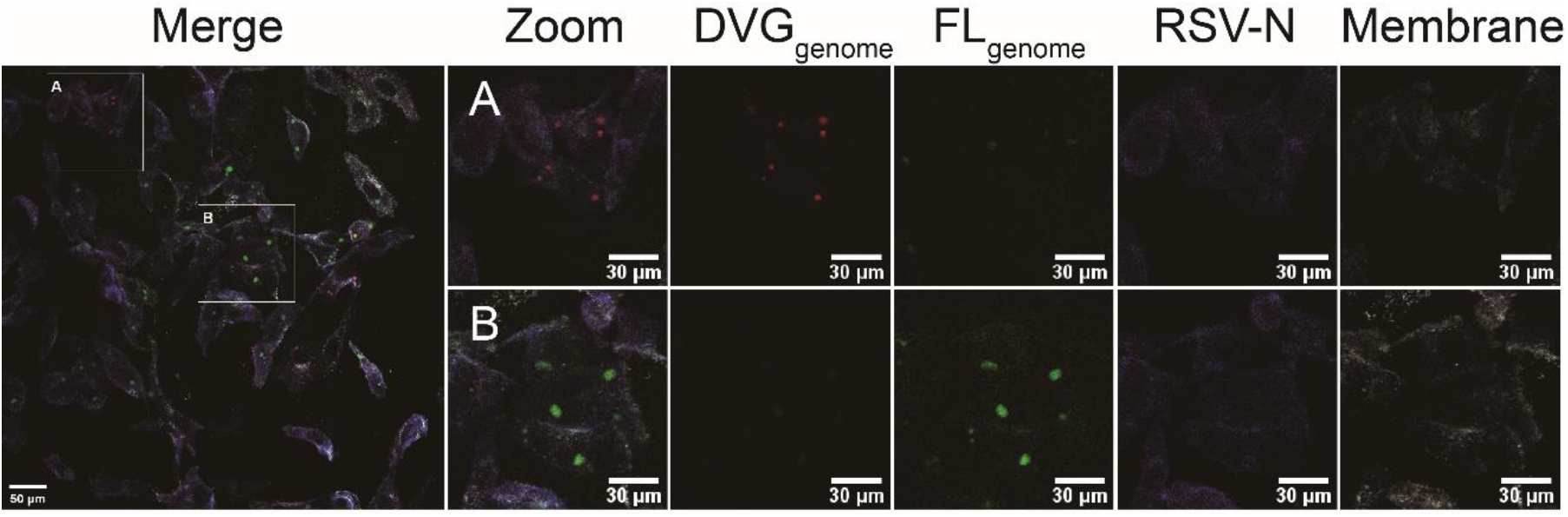
DVG-high cells gradually appeared in routine infection with RSV _low_DVG. RSV-A2 _low_DVG infected cells were co-stained with probes and antibody to examine the distribution of DVG_genome_ and Fl_genome_ and N protein. Cell-A was DVG-high, while cell-B was DVG-low.

